# Circular RNA profiling in the oocyte and cumulus cells reveals that *circARMC4* is essential for porcine oocyte maturation

**DOI:** 10.1101/586024

**Authors:** Zubing Cao, Di Gao, Tengteng Xu, Ling Zhang, Xu Tong, Dandan Zhang, Yiqing Wang, Wei Ning, Xin Qi, Yangyang Ma, Kaiyuan Ji, Tong Yu, Yunsheng Li, Yunhai Zhang

**Affiliations:** Anhui Province Key Laboratory of Local Livestock and Poultry, Genetical Resource Conservation and Breeding, College of Animal Science and Technology, Anhui Agricultural University, Hefei 230036, China

**Keywords:** circular RNA, Pig, Cumulus cells, Oocytes, Meiotic maturation

## Abstract

Thousands of circular RNAs (circRNAs) have been recently discovered in cumulus cells and oocytes from several species. However, the expression and function of circRNA during porcine oocyte meiotic maturation have been never examined. Here, we separately identified 7,067 and 637 circRNAs in both the cumulus cell and the oocyte via deep sequencing and bioinformatic analysis. Further analysis revealed that a faction of circRNAs is differentially expressed (DE) in a developmental stage-specific manner. The host genes of DE circRNAs are markedly enriched to multiple signaling pathways associated with cumulus cell function and oocyte maturation. Additionally, most DE circRNAs harbor several miRNA targets, suggesting that these DE circRNAs potentially act as miRNA sponge. Importantly, we found that maternal *circARMC4* knockdown by siRNA microinjection caused a severely impaired chromosome alignment, and significantly inhibited first polar body extrusion and early embryo development. Taken together, these results demonstrate for the first time that circRNAs are abundantly and dynamically expressed in a developmental stage-specific manner in cumulus cells and oocytes, and maternally expressed *circARMC4* is essential for porcine oocyte meiotic maturation and early embryo development.

## INTRODUCTION

Oocyte meiotic maturation is the last stage of oogenesis and is the indispensable prerequisite for fertilization, preimplantation development of the embryo and even term development (Conti and Franciosi, 2018). In mammals, oocytes meiotically arrested at the prophase I stage have to undergo germinal vesicle (GV) breakdown and subsequent first polar body extrusion to reach the metaphase stage of meiosis II (MII). Under the physiological conditions, the oocyte is typically enclosed in several layers of cumulus cells, thus its meiotic maturation is normally accompanying with the correct execution of cumulus cell function. Indeed, numerous studies showed that the reciprocal communications between the oocyte and its encircling cumulus cells are critical not only for the acquisition of both meiotic and developmental competence of the oocyte but also for the execution of cumulus cell function (Gilchrist et al., 2004; Gilchrist et al., 2008; Russell et al., 2016). Therefore, the identification of novel molecules expressed in either cumulus cells or oocytes could be informative in elucidating their relative contributions to oocyte meiotic maturation and development.

Pigs are increasingly being used as an ideal animal model for humans in reproductive medicine research because they share many similarities in the aspects of physiology (aneuploidy rate), developmental timing (oocyte maturation and early embryo development), and genetics with humans (Mordhorst and Prather, 2017). Although many efforts have been made to improve the oocyte maturation *in vitro* in pigs, its meiotic and developmental capacity is still suboptimal relative to that under the physiological states (Nagai, 2001; Yuan et al., 2017). This may be due to the imperfectness of *in vitro* culture conditions currently used that cannot achieve the optimal maturational outcomes, and there is inadequate information regarding the unique molecular mechanisms of porcine oocyte meiotic maturation (Sun and Nagai, 2003; Prather et al., 2009). It is known that oocyte maturation is intricately regulated by cumulus cell or oocyte itself derived non-coding RNAs, such as microRNA (miRNA) (Dallaire and Simard, 2016; Wright et al., 2016; Li et al., 2017b; Gay et al., 2018; Minogue et al., 2018), endogenous small interference RNA (siRNA) (Suh et al., 2010) and long non-coding RNA (lncRNA) (Taylor et al., 2015). Recently, circular RNA (circRNA) has received increasing attention in multiple biological research fields. CircRNA, formed by back-splicing of pre-mRNA transcripts, is a novel class of non-polyadenylated, single-stranded, covalently closed and long noncoding RNAs. Previous studies indicated that circRNA is mainly derived from exons, introns, intergenic regions (Lasda and Parker, 2014), and is widely distributed on the chromosomes in animal cells (Fan et al., 2015; Liang et al., 2017; Shen et al., 2019). CircRNA is discovered to frequently exert different molecular functions, such as transcriptional regulation, microRNA and RNA binding protein sponge, and mRNA trap (Bose and Ain, 2018). It is worthy noted that circRNA exhibits a higher resistance to degradation (Suzuki and Tsukahara, 2014) and is often expressed in a cell type- and developmental stage-specific manner (Fan et al., 2015; Dang et al., 2016). With respect to the field of animal reproduction, circRNA expression have been well characterized in different types of tissues and cells, including ovary (Liang et al., 2017; Cai et al., 2018; Chen et al., 2018), testis (Dong et al., 2016; Liang et al., 2017; Gao et al., 2018), placenta (Yan et al., 2018), follicle (Tao et al., 2017; Shen et al., 2019), spermatogenic cell (Lin et al., 2016), granulosa cell (Cheng et al., 2017; Fu et al., 2018), embryonic stem cell (ESC) (Yu et al., 2017), induced pluripotent stem cell (iPS) (Lei et al., 2018), germline stem cell (Li et al., 2017a), oocyte and embryo(Gardner et al., 2012; Fan et al., 2015; Dang et al., 2016). However, only two circRNAs in the reproduction field, namely *circBIRC6* and *circCORO1C*, have so far functionally been proved to be involved in the regulation of maintenance of ESC pluripotency and somatic cell reprogramming (Yu et al., 2017). Although the expression of circRNAs in porcine multiple tissues has been reported (Veno et al., 2015; Liang et al., 2017), its expression and function in porcine oocyte meiotic maturation remain unclear.

Here, we address the expression and function of circRNA in porcine oocyte meiotic maturation. Especially, we identified thousands of circRNAs in both the cumulus cell and the oocyte, some of which often display a developmental stage-specific expression. Unexpectedly, we found that maternally expressed *circARMC4* is an essential regulator for oocyte meiotic maturation and development. Therefore, our findings could have important implications in selecting potential biomarkers in cumulus cells to predict oocyte meiotic and developmental competence and developing strategies that improve assistant reproductive technique in humans.

## RESULTS

### Characterization of circRNAs expressed in porcine cumulus cell and oocyte during meiotic maturation

Since only the very limited amount of total RNA from thousands of porcine oocytes can be obtained, it is insufficient to meet the minimal needs of deep circRNA sequencing. To overcome the technical challenge, a mathematical method, named “complementary set function”, was applied to screen circRNAs expressed in oocytes during meiotic maturation. Thus, COCs and pure cumulus cells termed as GCC at GV stage, mixture samples of oocytes with first polar body extrusion and cumulus cells, and pure cumulus cells termed as MCC, were separately sequenced on the Illumina Hiseq platform (Fig. 1A). We then identified specific circRNAs expressed in oocytes by comparing circRNA transcripts of pure cumulus cells and mixture samples of cumulus cells and oocytes before and after maturation (Fig. 1A). To verify the reproducibility of sequencing data, we collected three sets of each sample. The correlation coefficient between two biological replicates ranged from 0.813 to 1, suggesting reliable sequencing data. A total of 803 million valid reads were obtained by removing the adapter and low-quality sequences and were mapped to the porcine reference genome (Table. S2). We totally identified 7,224 circRNAs from 3419 host genes, including 7,067 circRNAs from 3,329 host genes in the cumulus cell and 637 circRNAs from 476 host genes in the oocyte, respectively (Table. S3). The majority of host genes produce a single circRNA, whereas a few host genes generate multiple different circRNA species, specifically, 45.36% of host genes (1,510/3,329) in the cumulus cell and 19.12% (91/476) in the oocyte (Table. S4). The analysis of host gene distribution on chromosome revealed that circRNAs are widely transcribed from 18 autosomes and the X chromosome (Fig. 1B). Furthermore, chromosome 1 produces the most circRNAs in both the cumulus cell and the oocyte, whereas the least circRNAs are separately generated on chromosome 11 in the cumulus cell and chromosome 12 in the oocyte (Fig. 1B). The analysis of circRNA distribution in the genome indicated that most circRNAs is produced from exonic (60.82% in cumulus cell vs. 53.67% in oocyte) and intronic regions (28.45% vs. 32.99%), while a small fraction of circRNAs originate from intergenic regions (10.73% vs. 13.34%)(Fig. 1C). The average GC content of circRNA is around 47% in the cumulus cell and is approximately 45% in the oocyte, which are similar to those of linear mRNA molecules in pigs (Fig. 1D). Collectively, we identify and characterize thousands of circRNAs that are separately expressed in porcine cumulus cells and oocytes during meiotic maturation.

**Fig. 1.**
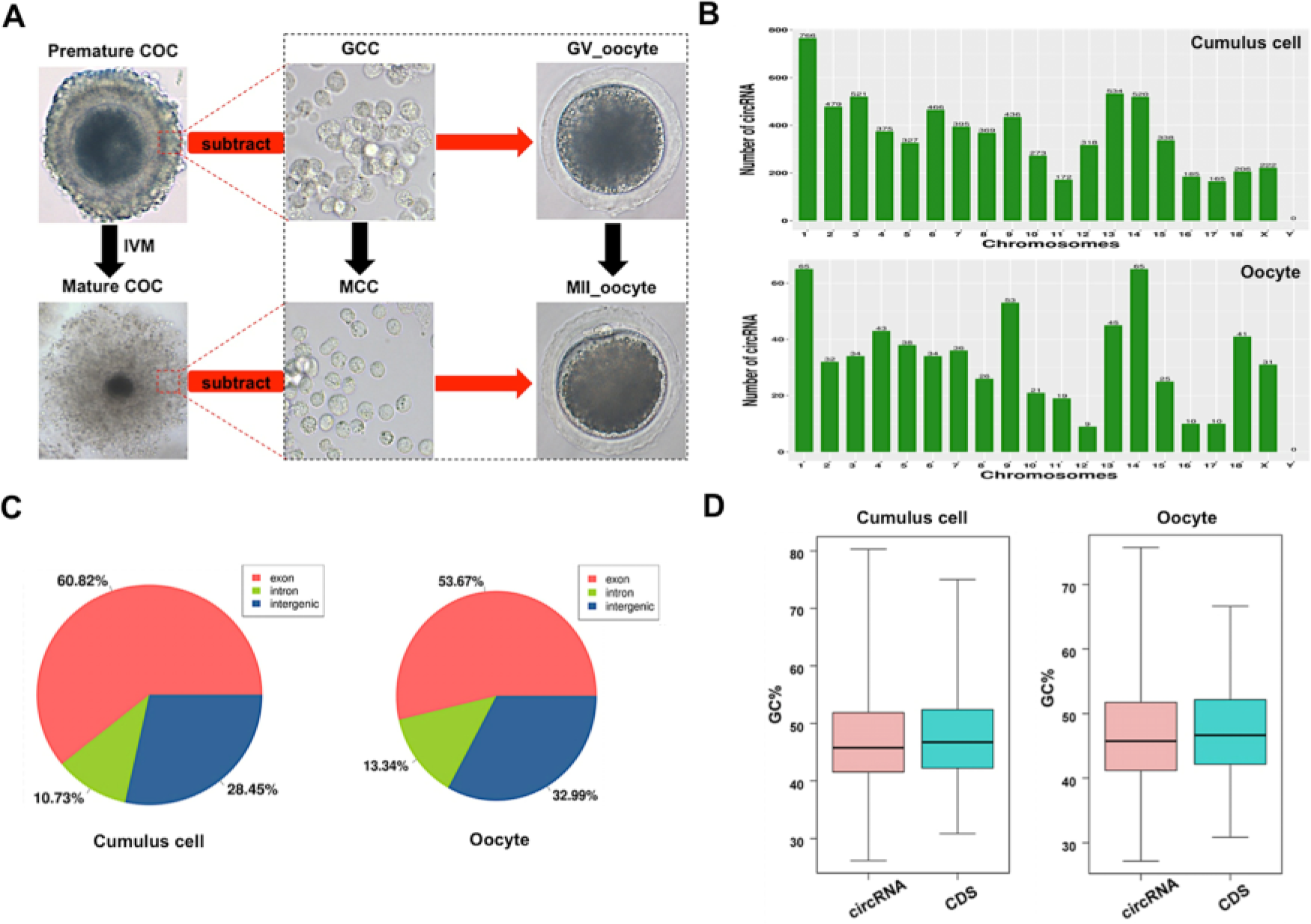
Characteristics of circRNAs expressed in porcine cumulus cells and oocytes. (A) Schematic illustration of experimental designs identifying circRNAs expressed in the cumulus cell or the oocyte before and after maturation. Premature COCs, cumulus cells before maturation (termed GCC), mature COCs, and cumulus cells after maturation (termed MCC) were collected respectively for RNA-seq. Of note, circRNAs expressed in the GCC were subtracted from these identical circRNAs expressed in the premature COCs to identify circRNAs expressed in GV oocytes, which are termed GV oocyte. Similarly, circRNAs expressed in MCC were subtracted from these identical circRNAs expressed in the mature COCs to identify circRNAs expressed in MII oocytes, which are termed MII oocyte. COCs, cumulus-oocyte complexes; GV, germinal vesicle; MII, metaphase II; IVM, in vitro maturation. Red dashed insets show cumulus cells before and after oocyte maturation at high magnification. (B) Chromosome distribution of circRNAs. Chromosome distribution of total circRNAs expressed in the cumulus cell or the oocyte was shown in the upper panel and bottom panel, respectively. (C) Genomic location of circRNAs. Genomic distribution of total circRNAs expressed in the cumulus cell or the oocyte was shown in the left panel and right panel, respectively. (D) GC enrichment of circRNAs and mRNAs. GC content of total circRNAs and mRNAs expressed in the cumulus cell or the oocyte was separately shown in the left panel and right panel.

### Identification and validation of differentially expressed circRNAs in both the cumulus cell and the oocyte

To evaluate dynamic changes of circRNA expression in both the cumulus cell and the oocyte during meiotic maturation, valid reads in RNA-sequencing data were quantified before and after oocyte maturation. Among all circRNAs detected in the cumulus cell, 77 circRNAs and 418 circRNAs are specifically expressed in GCC and MCC, respectively, whereas 6,572 circRNAs are commonly expressed in the cumulus cell between the two stages (Fig. 2A). Meanwhile, of all circRNAs identified in the oocyte, 428 circRNAs and 80 circRNAs are exclusively expressed in GV oocyte and MII oocyte, respectively, whereas 129 circRNAs are co-expressed in the oocyte between the two stages (Fig. 2A). In a comparison of MCC and GCC, the 928 host genes produce 1,902 differentially expressed circRNAs (DECs) (Table. S5) (*P* < 0.05), including 1,602 upregulated and 300 downregulated (Fig. 2B), while 191 circRNAs (9 upregulated and 182 downregulated) from 30 host genes are differentially expressed between MII oocyte and GV oocyte (Fig. 2B, Table. S5) (*P* < 0.05). Next, we performed hierarchical clustering analysis of the top 100 differentially expressed circRNAs in both the cumulus cell and the oocyte. As shown in the heatmap, samples at the same stages are clustered together, and the expression levels of circRNAs exhibit dynamic changes during oocyte maturation (Fig. 2C). To validate the circRNA sequencing data, the expression levels of 10 circRNAs (5 top upregulated in the cumulus cell and 5 top downregulated in the oocyte) before and after maturation, namely *circCORO1C*, *circVCAN*, *circLAPTM4B*, *circANXA2*, *circSCARB1* and *circZP4*, *circPRKCH*, *circCHL1*, *circVOCH1*, *circESRP1*, were analyzed by quantitative real-time PCR. These 10 circRNA candidates are first shown to be resistant to RNase R treatment, which verified their circularized characteristics (Fig. S2A, B, C, D). The expression patterns of these selected circRNAs during oocyte maturation are highly consistent with the treads obtained from circRNA sequencing data (Fig. 2D), confirming the results obtained by circRNA sequencing. Together, a fraction of circRNAs identified in both the cumulus cell and the oocyte exhibit stage-dependent dynamic expressions during meiotic maturation.

**Fig. 2.**
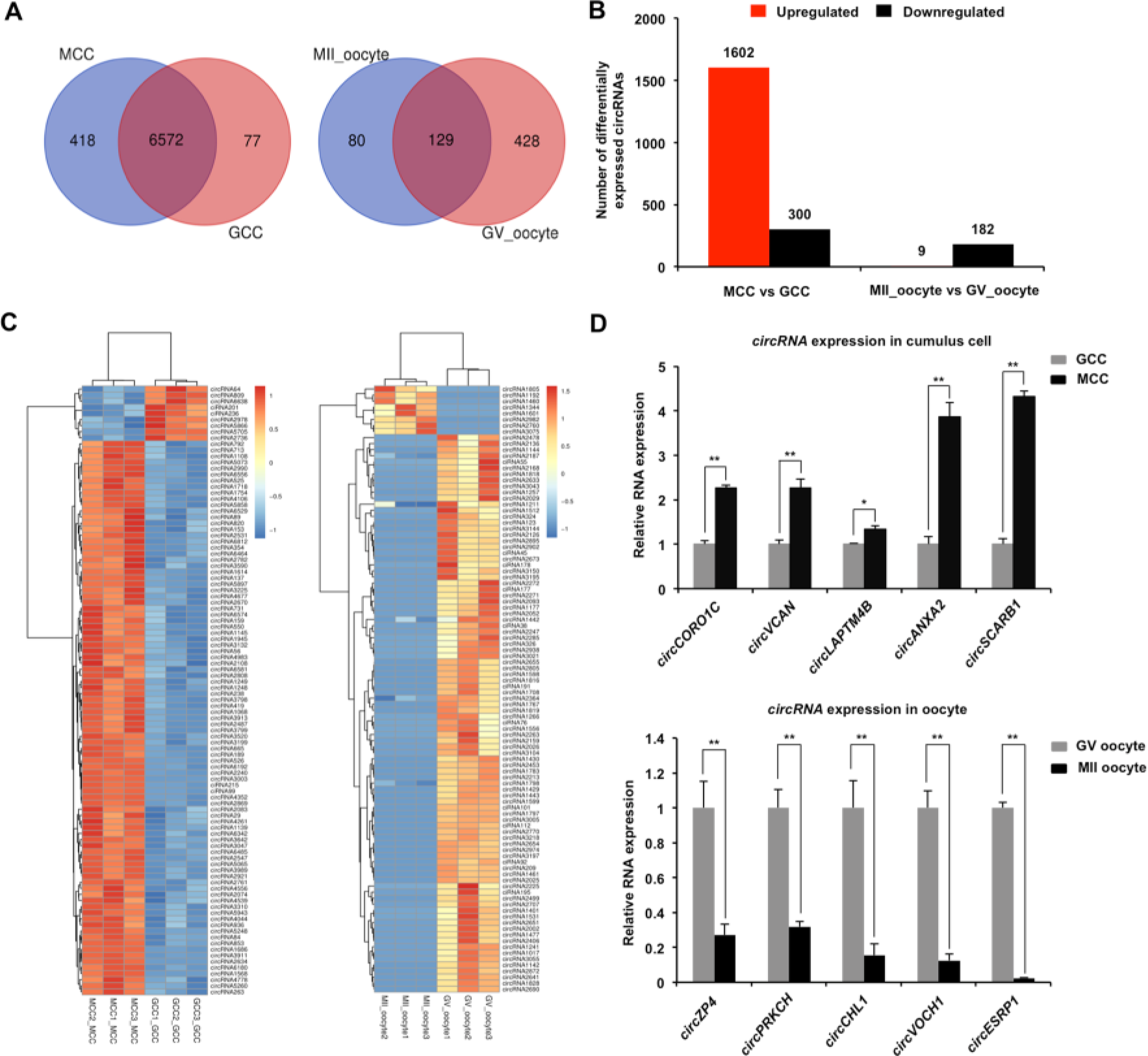
Identification and validation of differentially expressed circRNAs (DECs) in both the cumulus cell and the oocyte during meiotic maturation. (A) Venn diagram of circRNAs identified in the cumulus cell or the oocyte. Cumulus cells and oocytes before and after meiotic maturation were pooled for RNA-seq. Expression levels of circRNAs in the cumulus cell (left panel) and the oocyte (right panel) were analyzed by means of a binominal statistical test. Overlapping circles present circRNAs that are common for the cumulus cell or the oocyte between two different stages. Non-overlapping circles indicate circRNAs that are specific for the cumulus cell or the oocyte before (pink) and after (blue) meiotic maturation. (B) The number of differentially expressed circRNAs in the cumulus cell or the oocyte before and after meiotic maturation. The results were considered statistically significant at P_adjusted_< 0.05 and log2 fold change ≥1. Red bars indicate up-regulated circRNAs; green bars denote down-regulated circRNAs. (C) Heatmap illustrating the expression patterns of differentially expressed circRNAs in the cumulus cell (left panel) or the oocyte (right panel) before and after meiotic maturation. The red blocks represent up-regulated circRNAs, and the blue blocks represent down-regulated circRNAs. The color scale of the heatmap indicates the expression level, where the brightest blue stands for −1.0 log2 fold change and the brightest red stands for 1.0 or 1.5 log2 fold change. (D) Validation of the selected differentially expressed circRNAs identified in both the cumulus cell and the oocyte. The several selected circRNAs were chosen from top up and top down-regulated circRNAs in the cumulus cell or the oocyte. Relative abundance of circRNAs in the cumulus cell (upper panel) and the oocyte (bottom panel) was determined by qPCR. The data were normalized against endogenous housekeeping gene *EF1α1*, and the value for the cumulus cell or the oocyte at GV stage was set as one. The data are shown as mean ± S.E.M. Statistical analysis was performed using *t*-student test. Values with asterisks vary significantly, \**P* < 0.05, \*\**P* < 0.01.

### Functional analysis of host genes of differentially expressed circRNAs in both the cumulus cell and the oocyte

Under the assumption that circRNA functions may be relevant to the known functions of host genes, we performed GO and KEGG pathway analysis of the host genes producing DECs to predict their potential functions during oocyte maturation. The host genes generating DECs in both the cumulus cell and the oocyte were classified into three main categories (biological process, cellular component, and molecular function) according to the GO database. The 928 host genes producing DECs in the cumulus cell are totally enriched in 4,092 GO terms, among these, 146 GO terms are significantly enriched in three GO functions or undetermined function (*P* < 0.05), namely, 73 under “biological process”, 31 under “cellular component”, 41 under “molecular function” and 1 under “undetermined function” (Supplementary Table. S6). The top-ranking 25 biological processes, 15 cellular components, 10 molecular functions, and host genes involved in each GO term were listed (Fig. 3A, Table. S6), such as intracellular signal transduction (48 genes, e.g. *ADCY3*, *AKT3*, and *CDC42BPA*), negative regulation of cytoplasmic translation (4 genes, namely, *CPEB1*, *CPEB2*, *CPEB3*, and *CPEB4*), signal transduction by protein phosphorylation (12 genes, e.g. *BMPR1B*, *MAP3K2*, and *TGFBR1*), regulation of GTPase activity (14 genes, e.g. *ARHGAP6*, *PRKG1*, and *RICTOR*), cell-cell junction (19 genes, e.g. *ACTR3*, *AFDN*, and *CASK*). On the other hand, the 30 host genes producing DECs in the oocyte are totally enriched in 865 GO terms, of these, 236 GO terms are significantly enriched in three GO functions or undetermined function (Table. S6) (*P* < 0.05). The most significant biological processes, cellular components and molecular functions, and host genes involved in each GO term were listed (Fig. 3B, Table. S6) (*P* < 0.05), such as female meiotic division (2 genes, namely, *SYCP2* and *WEE2*), G-protein coupled receptor activity (3 genes, e.g. *CALCR*, *GABBR1*, and *SENP7*). In addition, the KEGG analysis of host genes producing DECs in the cumulus cell displayed 28 significant pathways (Fig. 3C, Table. S7) (*P* < 0.05), including progesterone-mediated oocyte maturation (16 genes), oocyte meiosis (17 genes), tight junction (17 genes), FoxO signaling (16 genes), MAPK signaling (21 genes), TGF-β signaling (10 genes), Wnt signaling (16 genes), Hippo signaling (14 genes). At the same time, we also found that the host genes generating DECs in the oocyte are enriched in 30 significant pathways (Fig. 3D, Table. S7) (*P* < 0.05), such as cAMP signaling (6 genes), cGMP-PKG signaling (5 genes), VEGF signaling (3 genes). Overall, the host genes-enriched aforementioned pathways in both the cumulus cell and the oocyte are mainly related to intercellular crosstalk between two cell types, or oocyte itself maturation. Therefore, these results indicated that DECs generated in both the cumulus cell and the oocyte probably exert critical functions in porcine oocyte maturation.

**Fig. 3.**
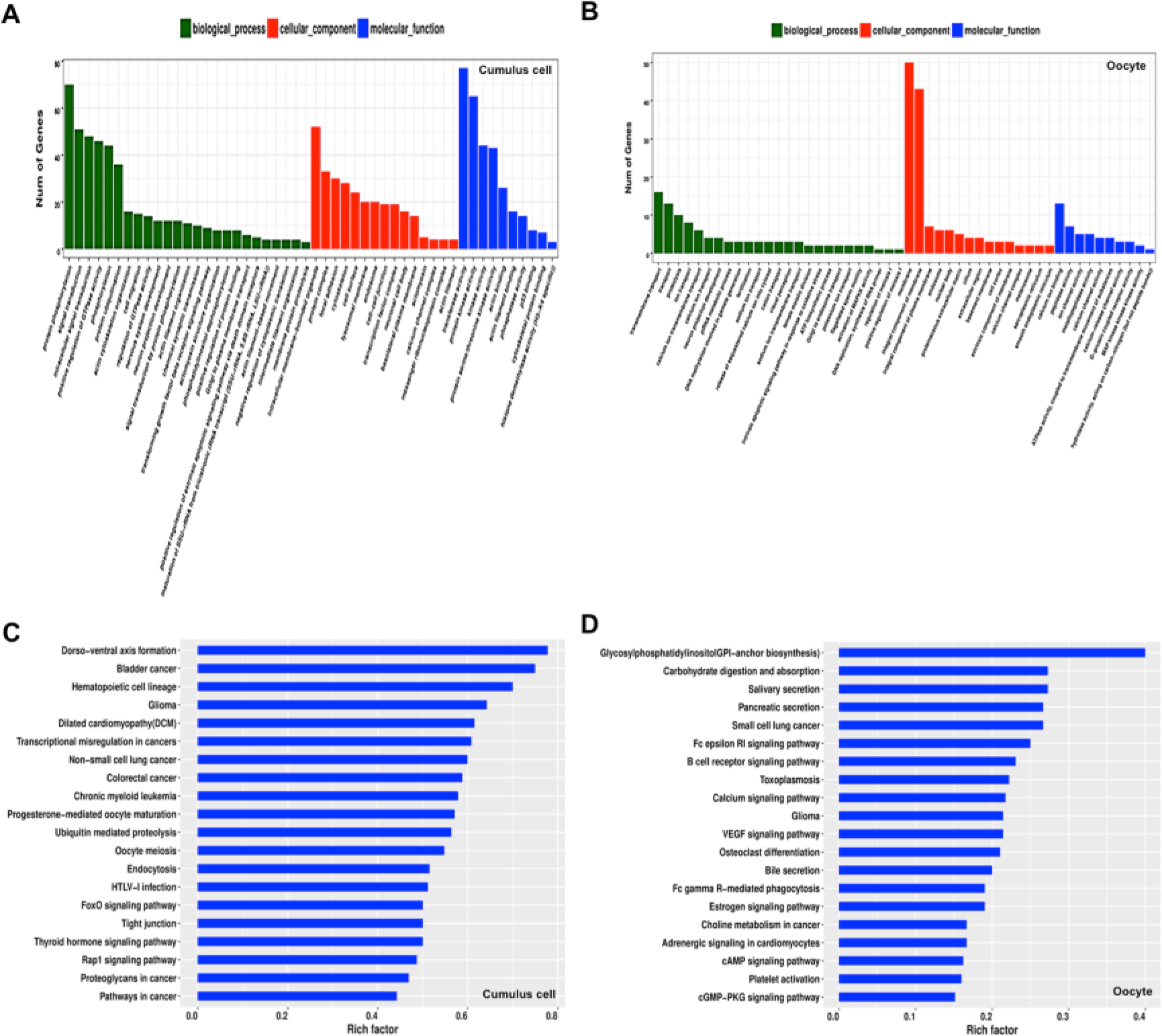
GO and KEGG analysis of host genes of differentially expressed circRNAs (DECs) in both the cumulus cell and the oocyte during meiotic maturation. (A) GO analysis of the top enriched terms of the differentially expressed circRNA hosting genes identified in the cumulus cell. Host genes of differentially expressed circRNAs were classified into three categories of the GO classification (blue bars: biological processes, green bars: cellular components and orange bars: molecular functions). (B) KEGG analysis of the top enriched signaling pathways of the differentially expressed circRNA hosting genes identified in the cumulus cell. (C) GO analysis of the top enriched terms of the differentially expressed circRNA hosting genes identified in the oocyte. Host genes of differentially expressed circRNAs were classified into three categories of the GO classification (blue bars: biological processes, green bars: cellular components and orange bars: molecular functions). (D) KEGG analysis of the top enriched signaling pathways of the differentially expressed circRNA hosting genes identified in the oocyte.

### Prediction of miRNA targets potentially sponged by circRNAs expressed in both the cumulus cell and the oocyte

It is previously reported that circRNA can sponge miRNAs to indirectly regulate gene expression in a post-transcriptional manner (Kulcheski et al., 2016). To assess whether all circRNAs identified in both the cumulus cell and the oocyte function as miRNA sponge, we predicted miRNA targets of these circRNAs using bioinformatic tools. We found that 7,165 out of 7,224 circRNAs (99.18%) have miRNA binding sites, whereas a very few circRNAs are predicted to have no potential miRNA targets. Furthermore, 411 putative microRNA targets on 7,165 circRNAs were commonly predicted using Targetscan and miRanda (Fig. 4A). Each circRNA may bind to one or multiple miRNA targets. Thus, the proportion of circRNA containing different numbers of miRNA targets is further measured in our study. Most circRNAs have at least two miRNA binding sites, of which the proportion of circRNA containing 6-10 miRNA targets is the highest one (Fig. 4B). Subsequently, we found 411 and 373 putative miRNA targets for 1,897 out of 1,902 DECs in the cumulus cell and 180 out of 191 DECs in the oocyte, respectively. The interaction relationships between selected representative DECs in both the cumulus cell and the oocyte and their predicted miRNA targets are shown (Fig. 4C, D). These data revealed that most DECs in the two cell types indeed contain multiple miRNA targets. It is noted that some miRNAs sponged by circRNAs have been shown to control cumulus cell function, oocyte maturation, and early embryo development. For example, miRNA-224 targeted by *circRNA3798* in cumulus cells (Li et al., 2017b), miRNA-21 bound by *circRNA193* (Wright et al., 2016), and miRNA-378 bound by *circRNA2982* (Pan et al., 2015) in oocytes. Altogether, these results demonstrate that circRNAs identified in both porcine cumulus cells and oocytes could have similar effects of miRNA sponge as those observed in other cellular contexts.

**Fig. 4.**
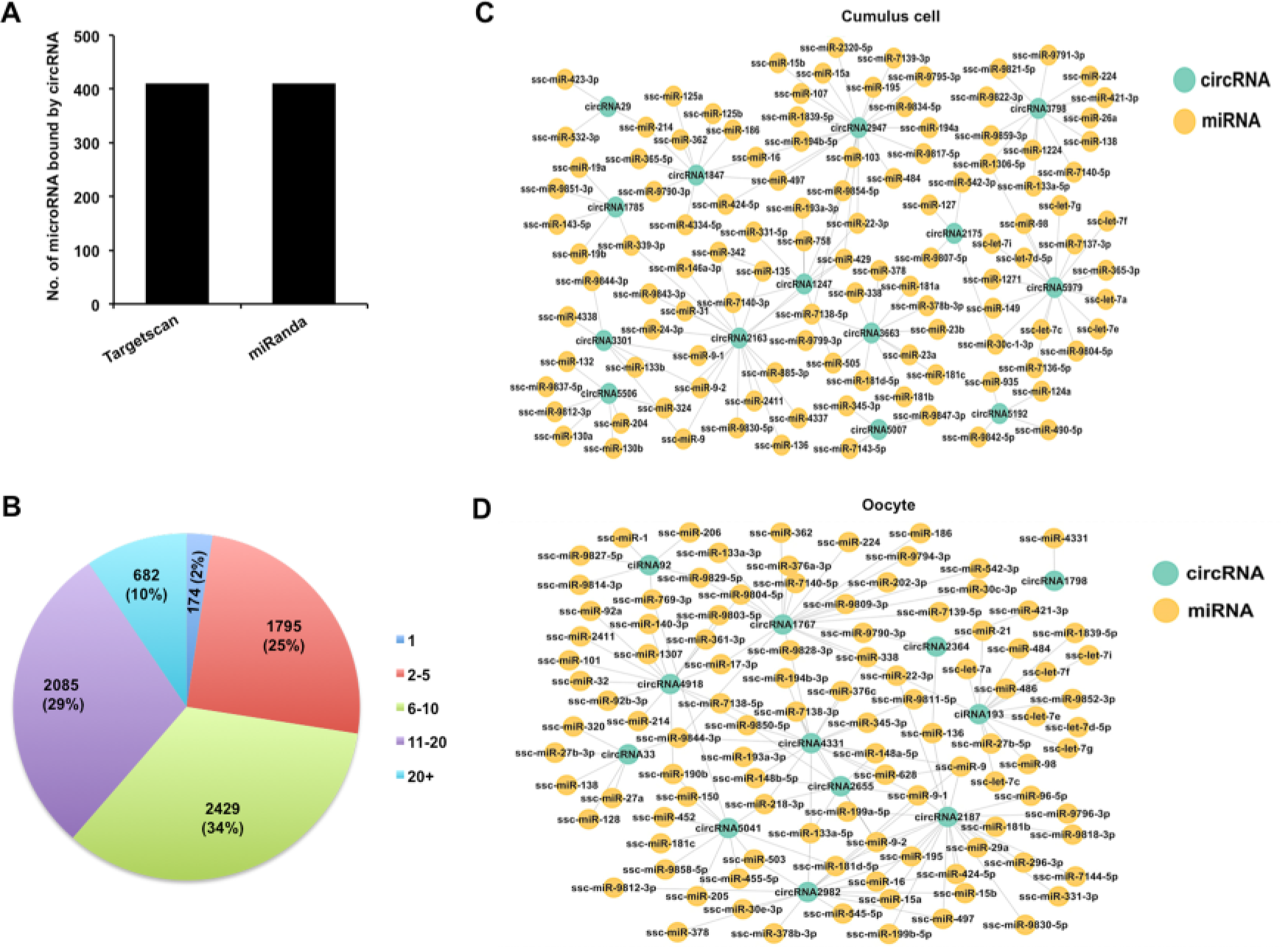
Analysis of interaction between DECs and miRNAs in both the cumulus cell and the oocyte. (A) Analysis of number of miRNAs for circRNAs by Targetscan and miRanda. (B) Analysis of the proportion of circRNA processing different numbers of miRNA targets. (C) Analysis for predicted targeted miRNAs of the selected DECs identified in the cumulus cell. The selected circRNAs were chosen from top DECs in the cumulus cell. Blue circles represent circRNA, and yellow circles represent miRNA. (D) Analysis for predicted targeted miRNAs of the selected DECs identified in the oocyte. The selected circRNAs were chosen from top DECs in the oocyte. Blue circles represent circRNA, and yellow circles represent miRNA.

### *CircARMC4* knockdown impairs porcine oocyte meiotic maturation and chromosome alignment

To further determine whether circRNAs identified in porcine oocytes by RNA sequencing function in meiotic maturation, one top upregulated circRNA, namely, *circRNA2982* (also called as *circARMC4*) from the host gene *ARMC4*, was selected for functional research. Firstly, the relative abundance of *circARMC4* in oocytes before and after maturation was analyzed by qPCR. We found that the expression levels of *circARMC4* in MII oocytes are significantly higher than those in GV oocytes (Fig. 5A), which is consistent with the tread obtained from RNA sequencing data. Furthermore, *circARMC4* transcripts, but not its corresponding linear counterparts, are resistant to RNase R treatment, which confirmed its circularized feature (Fig. S2C, D). Sanger sequencing of qPCR products spanning back-splicing sites between exon 14 and exon 15 also verified that it is indeed real circRNA (Fig. 5B). To examine the role of *circARMC4* in oocyte maturation, *circARMC4* was knocked down by microinjecting siRNA into GV oocytes. Results revealed that siRNA could significantly reduce the expression of *circARMC4*, but not its linear counterpart (Fig. 5C, D). Phenotypically, we found that *circARMC4* knockdown could apparently reduce the rate of first polar body extrusion (Fig. 5E, F). Also, oocytes with first polar body in control groups display bipolar spindles and normal linear chromosome morphology, while chromosomes in the substantial fraction of *circARMC4* knocked down oocytes are misaligned at the metaphase plate even though the spindles are morphologically normal (Fig. 5G). We further observed that the proportion of MII oocytes with abnormal chromosomes in the *circARMC4* knockdown group is significantly higher than that in control groups (Fig. 5H). Therefore, these data document that *circARMC4* is essential for oocyte meiotic maturation in pigs.

**Fig. 5.**
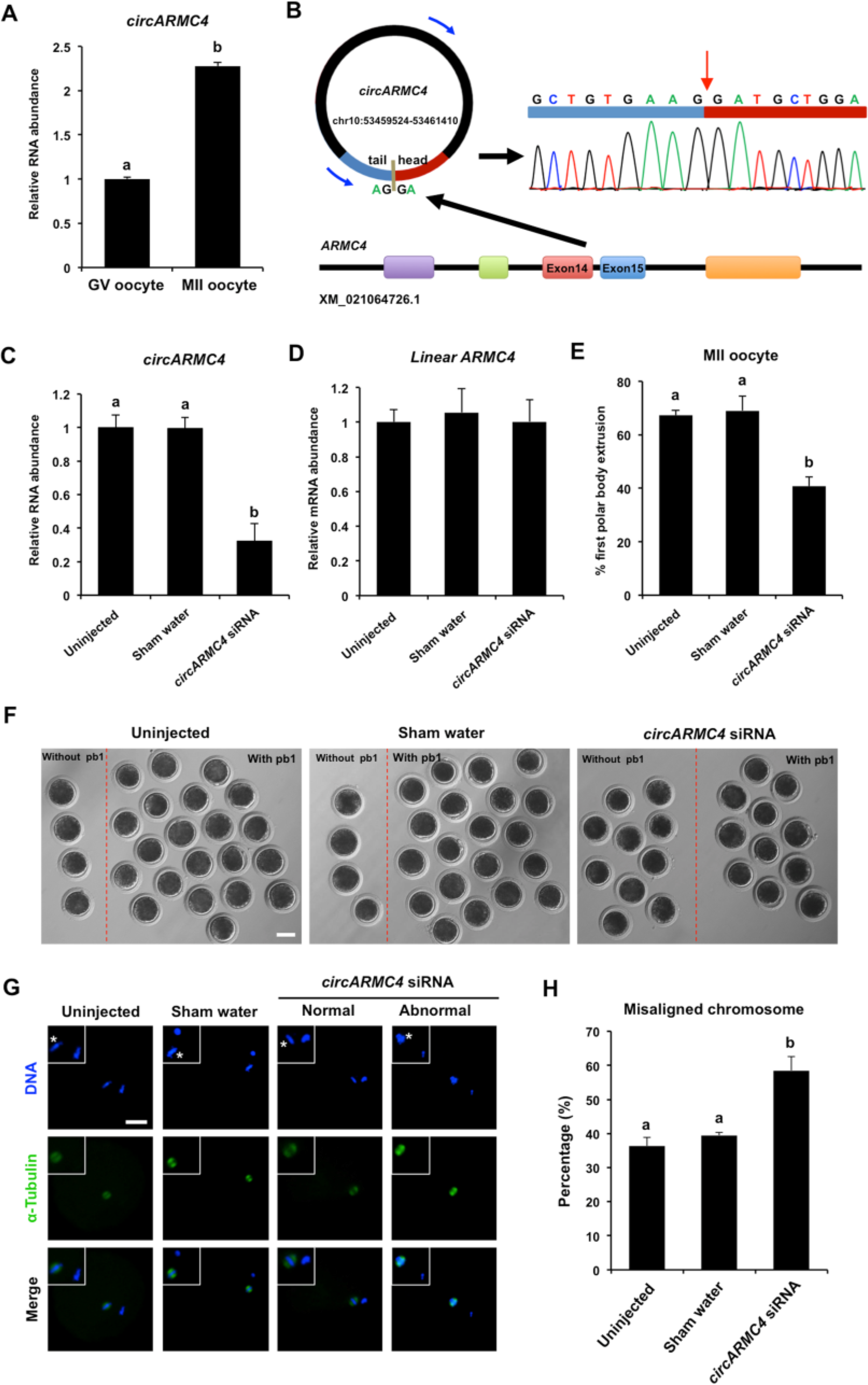
Effect of *circARMC4* knockdown on oocyte meiotic maturation and chromosome alignment. (A) *CircARMC4* expression in both the GV oocyte and the MII oocyte. Relative expression of *circARMC4* was determined by qPCR. The data are analyzed using student’s *t* test and are shown as mean ± S.E.M. Different letters on the bars indicate significant differences (*P* < 0.05). (B) Schematic illustration showed the ARMC4 exon 14 and exon 15 circularization forming circARMC4 (blue arrow). The presence of *circARMC*4 was validated by qPCR, followed by Sanger sequencing. Red arrow represents “head-to-tail” *circARMC4* splicing sites. The expression levels of *circARMC4* (C) and linear *ARMC4* (D) in the MII oocytes derived from GV oocytes. GV oocytes were injected with circARMC4 siRNA, followed by maturation *in vitro* for 44 h. Oocytes injected with water and uninjected oocytes were served as a sham control and a blank control, respectively. One hundred matured oocytes were collected for qPCR analysis. Relative abundance of *circARMC4* and *linear ARMC4* was determined by qPCR from four independent replicates. The data were normalized against endogenous housekeeping gene *EF1α1* and the value for the blank control was set as one. The data are shown as mean ± S.E.M. One-way ANOVA was used to analyze the data and different letters on the bars indicate significant differences (*P* < 0.05). (E) Analysis of the rate of oocyte maturation. The number of oocytes with first polar body after *in vitro* maturation for 44 h was recorded and the rate of first polar body extrusion was statistically analyzed by one-way ANOVA. The experiment was repeated four times with at least 100 oocytes per group. The data are shown as mean ± S.E.M and different letters on the bars indicate significant differences (*P* < 0.05). (F) Representative images of oocytes after *in vitro* maturation. The oocytes without pb1 and the oocytes with pb1 were shown in both the left side and the right side of each image, respectively. Scale bar: 100 μm. (G) Spindle and chromosome morphology in MII oocytes. Matured oocytes were stained for α-tubulin (green) and DAPI (blue). Shown are representative images obtained using confocal laser-scanning microscopy. The experiment was independently repeated three times with at least 40 oocytes per group. Bottom panel in each group shows the merged images between α-tubulin and DNA. White square insets indicate both spindles and chromosomes at high magnification. Asterisks indicate chromosomes. Scale bar: 50 μm. (H) Analysis of the percentage of oocytes with abnormal chromosome morphology. The chromosome morphology of MII oocytes was scored according to a published standard method. The data were statistically analyzed by one-way ANOVA. The data are shown as mean ± S.E.M and different superscripts on the bars indicate significant differences (*P* < 0.05).

### *CircARMC4* knockdown reduces developmental competence of porcine early embryos

Given that developmental competence of embryos largely depends on oocyte quality (Conti and Franciosi, 2018), we thus attempted to explore whether *circARMC4* knockdown in oocytes affects early development of porcine embryos. MII oocytes from both *circARMC4* knockdown group and two control groups were parthenogenetically activated and cultured up to the blastocyst stage. We found that *circARMC4* knockdown significantly reduced developmental efficiency of 2-cell, 4-cell, 8-cell embryos and blastocysts (Fig. 6A, B), suggesting that *circARMC4* knockdown could impair the developmental competence of oocytes. Hence, these results indicate that *circARMC4* knocked down oocytes possess poor developmental competence.

**Fig. 6.**
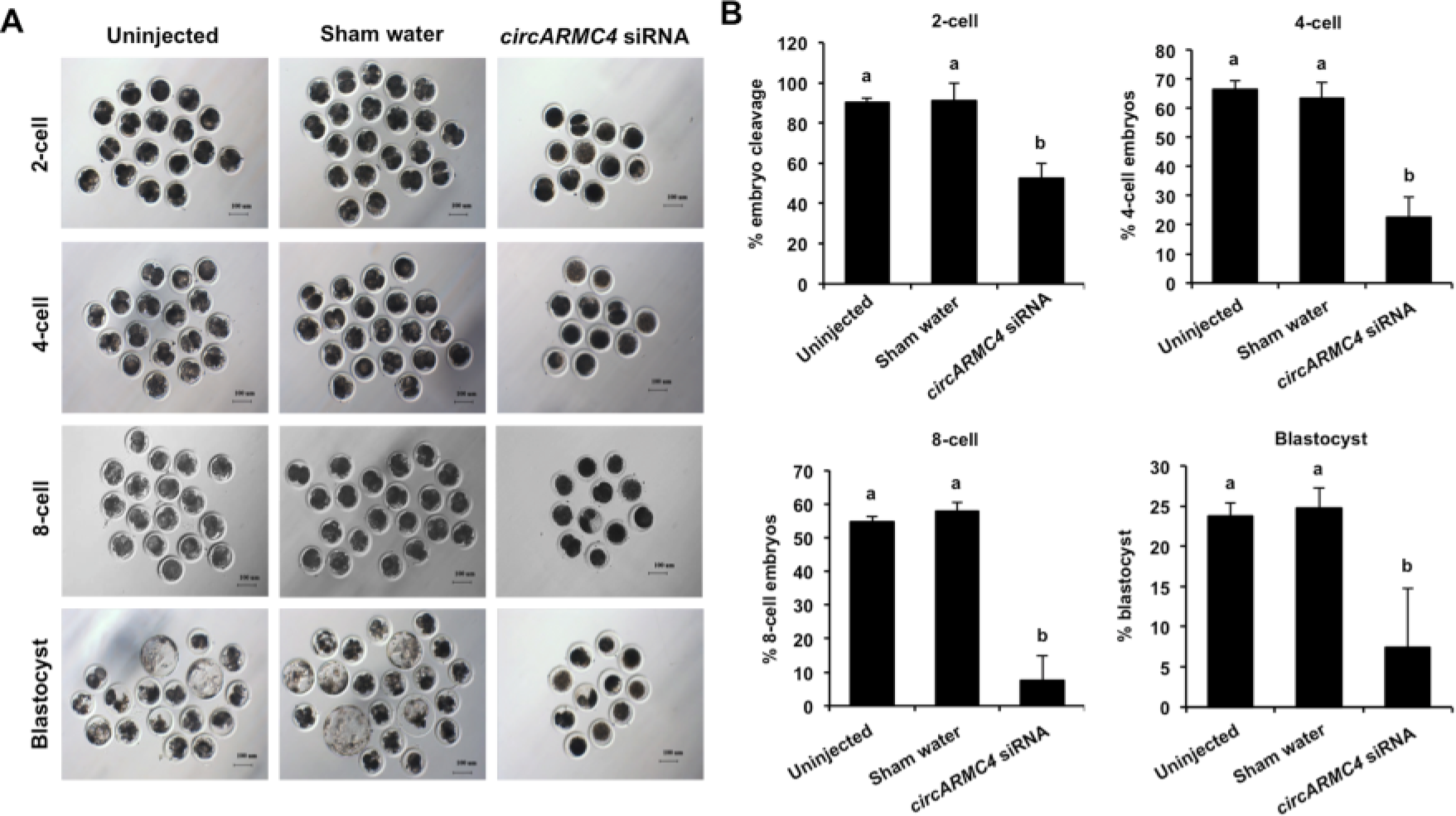
Effect of *circARMC4* knockdown on developmental competence of porcine early embryos. (A) Representative brightfield images of embryos at different developmental stages. GV oocytes were injected with siRNA or water and GV oocytes without any treatment, followed by maturation *in vitro* for 44 h. Oocytes with first polar body were parthenogenetically activated and cultured up to the blastocyst stage. The brightfield images of 2-cell, 4-cell, 8-cell embryo and blastocyst were captured by epifluorescence microscopy. Scale bar: 100 μm. (B) Analysis of the developmental rate of embryos at different developmental stages. The number of embryos at different developmental stages was recorded and the corresponding data were statistically analyzed by one-way ANOVA. All experiment was repeated four times. The data are shown as mean ± S.E.M and different letters on the bars indicate significant differences (*P* < 0.05).

## DISCUSSION

Although the expression profiles of circRNAs in different porcine tissues have been well studied (Veno et al., 2015; Liang et al., 2017), its expression and function in porcine oocyte meiotic maturation are yet to be characterized. We here identified thousands of circRNAs expressed in both the cumulus cell and the oocyte by deep RNA sequencing, some of which show spatio-temporal specific differential expression during meiotic maturation. Functional annotations of these DECs revealed that they could be engaged in the regulation of oocyte meiotic maturation through sponging microRNAs. Importantly, we further showed that *circARMC4*, a top up-regulated circRNA in the oocyte, is required for porcine oocyte meiotic maturation and early embryo development. Therefore, these results indicate that circRNAs expressed in both the cumulus cell and the oocyte could play a critical role in porcine oocyte meiotic maturation and early embryo development.

It is well known that cumulus cell potentiates oocyte meiotic and developmental competence after fertilization (Dumesic et al., 2015); conversely, the oocyte also promotes the proliferation and differentiation of cumulus cells during meiotic maturation (Gilchrist et al., 2008). Thus, identifying unique molecules expressed in either cumulus cells or oocytes could lay a foundation for elucidating its roles in each cell type. In this study, we detected 7,067 circRNAs in the cumulus cell and 637 circRNAs in the oocyte, respectively, indicating that cell type-dependent characteristics of circRNA expression and differential regulation of circularization events in different cells. A similar phenomenon is also observed in other studies (Liang et al., 2017; Vo et al., 2019). In addition, it is evident that cumulus cell is transcriptionally high active in the process of oocyte maturation while oocyte always retains in a transcriptionally quiescent state (Reyes and Ross, 2016). Cumulus cell could produce much more transcripts relative to the stored maternal transcripts in the oocyte, and circRNA biogenesis mainly depends on the back-splicing of pre-mRNA transcribed in the cell (Lasda and Parker, 2014). We thus speculated that a larger amount of circRNAs produced in the cumulus cell could be attributed to its higher transcriptional activity. Interestingly, we noted that the number of circRNAs identified in the cumulus cell or the oocyte apparently differs from that found in other species, such as cattle (Sun and Nagai, 2003), Xenopus (Gardner et al., 2012), mice (Fan et al., 2015) and humans (Dang et al., 2016), suggesting that circRNA expression may have a feature of species diversity. We also found that some of the host genes produce multiple different circRNA species in our analyzed samples, which could be explained by the fact that back-splicing of pre-mRNA transcripts from the same host gene frequently occur between different genomic sites (Lasda and Parker, 2014). CircRNAs detected in this study are widely distributed on the 18 autosomes and the X chromosome, indicating functional diversity of circRNAs. Meanwhile, the number of circRNAs originated from each chromosome in both the cumulus cell and the oocyte also varies, which may be associated with the chromosome length. Regarding the origin of circRNAs in both the cumulus cell and the oocyte, it was shown that circRNAs mainly originate from exons, which is consistent with the previous studies (Liang et al., 2017). Of note, exon-derived circRNAs in the other cellular contexts are preferentially involved in the post-transcriptional regulation of gene expression (Bose and Ain, 2018), which is also likely adaptable to cumulus cell and oocyte. Furthermore, consistent with data reported in other porcine cell types (Liang et al., 2017), the GC content of circRNA in both the cumulus cell and the oocyte is similar to that of linear mRNA molecules, implying that thermal stability of circRNA could be conserved across different cell types. Based on the above analysis, it is concluded that circRNAs identified in our study harbor some unique or common features between different cell types or species.

Dynamic changes of gene expression patterns usually reflect their potential functions within specific cells at concrete time points during development. Previous studies have revealed that circRNAs exhibit tissue- and developmental stage-specific expression in pigs (Liang et al., 2017), mice (Fan et al., 2015) and humans (Dang et al., 2016). Likewise, we also found that circRNAs are expressed in a stage-specific manner during oocyte maturation. Specifically, the number of circRNAs gradually increases in the cumulus cell that has a high proliferation activity, suggesting that circRNA expression could be positively correlated to the proliferation activity of cumulus cells, which is completely contrary to that observed in cancer cells (Bachmayr-Heyda et al., 2015). This discrepancy could be due to the differences in cell types or cellular physiological states. On the other hand, the number of circRNAs progressively decreases in the oocyte with the quiescent transcriptional state. It is thus speculated that the reduction of circRNA expression may be associated with the intrinsic active degradation machinery of maternal transcripts in oocytes. Furthermore, some of the circRNAs identified in both the cumulus cell and the oocyte are differentially expressed between GV and MII stages, implying that the fluctuation of circRNA abundance seems likely to be related to their specific roles in porcine oocyte maturation.

Potential functions of circRNAs in porcine cumulus cells and oocytes can be indirectly predicted by analyzing their host genes. GO and KEGG analysis of host genes producing DECs in cumulus cells revealed that GO terms are mainly enriched in the signal transduction related biological processes and pathways, such as regulation of GTPase activity and tight junction. The GTPase signaling pathway is reported to play critical roles in a wide range of cellular processes (Oliveira and Yasuda, 2014). Previous studies indicated the activation of GTPases is responsible for the accumulation of cell junction proteins to regulate the establishment of cell junctions and cell-cell adhesion between oocytes and cumulus cells (Fukata and Kaibuchi, 2001; Jiang et al., 2017). The GTPase signaling pathway is also involved in controlling the proliferation and differentiation of chicken granulosa cells (Shen et al., 2019). Tight junction is shown to regulate the transport of macromolecules from granulosa cells to oocytes in chicken (Schuster et al., 2004). Based on these studies and our results, it is plausible that DECs identified in the cumulus cell may be engaged in the intercellular communications between porcine oocytes and cumulus cells. Moreover, other signaling pathways related to animal reproduction, eg. Progesterone-mediated oocyte maturation, FoxO signaling, MAPK signaling, TGF-β signaling, Wnt signaling, and Hippo signaling, are also significantly enriched. Especially, small GTPase RhoA (Zhang et al., 2014), MAPK signaling members (Li et al., 2008) and TGF-β family protein GDF8 (Yoon et al., 2017) are essential for porcine oocyte maturation while Wnt signaling pathway negatively regulates porcine oocyte maturation (Shi et al., 2018). On the other hand, GO terms significantly enriched by host genes generating DECs in oocytes are mainly related to oocyte meiosis, such as female meiotic division, cAMP signaling, and VEGF signaling. For instance, both WEE2 enriched in the female meiotic division and cAMP molecules have been shown to inhibit the resumption of oocyte meiosis in several species (Hanna et al., 2010; Ramos Leal et al., 2018). In contrast, VEGF supplement significantly improved maturation rate of oocytes in cattle (Luo et al., 2002), sheep (Yan et al., 2012) and pigs (Bui et al., 2017). Altogether, we thus reasoned that DECs identified in the oocyte could positively or negatively regulate porcine oocyte meiotic maturation in a synergistic manner.

It has been shown that exon-derived circRNAs preferentially function in the post-transcriptional regulation (Bose and Ain, 2018). Similarly, we found that the vast majority of DECs identified in this study not only stem from the exonic sequence but also have putative miRNA binding site, which indicates that most DECs probably act as miRNA sponge. We observed that the number of miRNAs bound by each circRNA varies, which may be related to circRNA length and the inherent features of circRNA sequences. CircRNA-miRNA network analysis further revealed that DECs in the analyzed samples can interact with miRNA at multiple direction levels, which is consistent with that observed in the previous studies (Liang et al., 2017; Shen et al., 2019). This potential interplay between circRNA and miRNA could provide a reference for elucidating regulatory mechanisms of these DECs in oocyte meiotic maturation. Indeed, recent studies have shown that maternal miRNA participates in regulating oocyte maturation and early embryo development in several species, including pigs (Wright et al., 2016), medaka (Gay et al., 2018), and *C.elegans* (Minogue et al., 2018). Specifically, miRNA-7 was found to inhibit epidermal growth factor receptor (EGFR) expression in human cancer cells (Webster et al., 2009) and EGFR signaling is required for cumulus cell expansion and oocyte maturation (Prochazka et al., 2017), suggesting that miRNA-7 may negatively regulate oocyte maturation. Maternal miRNA-21 (Wright et al., 2016) or cumulus cell-derived miRNA-224 (Li et al., 2017b) and miRNA-378 (Pan et al., 2015) positively or negatively regulate porcine oocyte maturation and early embryo development. Our results indicated that these abovementioned miRNAs could be bound by DECs expressed in both the cumulus cell and the oocyte. Therefore, these DECs may block the functional roles of miRNAs by sequestering them, thereby promoting or inhibiting porcine oocyte maturation and early embryo development.

Examining the direct roles of individual circRNAs in oocyte meiotic maturation and embryo development should be subject in future investigations. In the current study, a maternally expressed *circARMC4* was selected to explore its roles in porcine oocyte maturation. We discovered that *circARMC4* knockdown led to a significant reduction in the rate of oocyte maturation and early embryo development and a higher proportion of misaligned chromosome. To our knowledge, this is proved for the first time that circRNA is essential for oocyte maturation and early embryo development. It is previously reported that Gudu is the Drosophila homolog of mammalian *ARMC4* which is the host gene of *circARMC4* and has been shown to be required for spermatogenesis, but not female fertility (Cheng et al., 2013), predicting that *circARMC4* could be involved in the regulation of mammalian reproduction. Besides, miRNA-378 bound by *circARMC4* is a negative regulator of porcine oocyte maturation and early embryo development (Pan et al., 2015). It is thus possible that the above-observed phenotypes may be caused by *circARMC4* knockdown-mediated activation of miRNA-378. Of note, circRNA also can execute roles in other models of post-transcriptional regulation. We cannot thus exclude the possibility that *circARMC4* might exert functions in porcine oocyte maturation and early embryo development via other mechanisms, such as RNA binding protein sponge, mRNA trap and circRNA itself translation. These hypotheses, however, need to be further established by additional experimental designs in future studies.

In conclusion, these results demonstrate that cumulus cells and oocytes generate abundant circRNAs during meiotic maturation, of which thousands of circRNAs are expressed in a developmental stage-specific manner. Our data also document for the first time that maternally expressed *circARMC4* is essential for porcine oocyte meiotic maturation and early embryo development.

## MATERIALS AND METHODS

### Ethics statement

All experiments using pigs were carried out according to the guidelines of the Institutional Animal Care and Use Committee (IACUC) at Anhui Agricultural University.

### *In vitro* maturation of oocytes

Ovaries were collected from a local slaughterhouse and transported to the laboratory at 28-35 °C in physiological saline solution. Follicular fluids from antral follicles of different diameters at 1-2 mm, 3-6 mm, and more than 6 mm were aspirated using a sterile 10 ml syringe with 18 gauge needles. Cumulus-oocyte complexes (COCs) from each type of follicles were then selected under a stereomicroscope. Subsequently, appropriately 50 COCs were transferred to 400 μl *in vitro* maturation medium (TCM-199 supplemented with 5% FBS, 10% porcine follicular fluid, 10 IU/ml eCG, 5 IU/ml hCG, 100 ng/ml L-Cysteine, 10 ng/ml EGF, 100 U/ml penicillin and 100 mg/ml streptomycin) covered with mineral oil in 4-well plate and cultured for 44 h at 38.5 °C, 5% CO_2_, 95% air and 100% humidity. Cumulus cells surrounding oocyte were removed by gentle pipetting in 1 mg/ml hyaluronidase solution. The normal nuclear maturation of oocytes was indicated by first polar body (pb1) extrusion.

### Collection of RNA sequencing samples

Four types of sample were collected to meet the minimum amount of total RNA for RNA sequencing. First, the mixed cells consisting of fully-grown GV oocytes and cumulus cells were collected. Second, the cumulus cells isolated from fully-grown cumulus-oocyte complexes were solely collected and were also termed GCC. Third, after meiotic maturation of oocytes, the mixed cells containing MII oocytes and its surrounding cumulus cells were collected. Fourth, the cumulus cells encircling MII oocyte were only collected and were also termed MCC.

### circRNA library construction and sequencing

Total RNA was isolated and purified using Trizol reagent (Invitrogen, Carlsbad, CA, USA) following the manufacturer’s procedure. The RNA amount and purity of each sample was quantified using NanoDrop ND-1000 (Wilmington, DE, USA). The RNA integrity was assessed by Agilent 2100 with RIN number >7.0. Approximately 5 ug of total RNA was used to deplete ribosomal RNA according to the manuscript of the Ribo-Zero™ rRNA Removal Kit (Illumina, San Diego, USA). After removing ribosomal RNAs, the left RNAs were fragmented into small pieces using divalent cations under high temperature. Then the cleaved RNA fragments were reverse-transcribed to create the cDNA, which was next used to synthesize U-labeled second-stranded DNAs with E. coli DNA polymerase I, RNase H and dUTP. An A-base is then added to the blunt ends of each strand, preparing them for ligation to the indexed adapters. Each adapter contains a T-base overhang for ligating the adapter to the A-tailed fragmented DNA. Single-or dual-index adapters are ligated to the fragments, and size selection was performed with AMPureXP beads. After the heat-labile UDG enzyme treatment of the U-labeled second-stranded DNAs, the ligated products are amplified with PCR by the following conditions: initial denaturation at 95 °C for 3 min; 8 cycles of denaturation at 98 °C for 15 sec, annealing at 60 °C for 15 sec, and extension at 72 °C for 30 sec; and the final extension at 72 °C for 5 min. The average insert size for the final cDNA library was 300 bp (±50 bp). At last, we performed the paired-end sequencing on an Illumina Hiseq 4000 (LC Bio, China) following the vendor’s recommended protocol.

### Bioinformatic analysis of circRNA

Firstly, Cutadapt was used to remove the reads that contained adaptor contamination, low quality bases and undetermined bases. Then sequence quality was verified using FastQC. We used Bowtie2 and Tophat2 to map reads to the genome of Sus scrofa (Ensemble database). Remaining reads (unmapped reads) were still mapped to the genome using Tophat-fusion. CIRCExplorer was used to de novo assemble the mapped reads to circular RNAs at first; Then, back splicing reads were identified in unmapped reads by tophat-fusion and CIRCExplorer. All samples were generated unique circular RNAs. The differentially expressed circRNAs were selected with log2 (fold change) > 1 or log2 (fold change) < −1 and with statistical significance (p value < 0.05) by t-test. To analyze functions of differentially expressed circRNAs and their host genes of circRNAs involvement in the common biological processes, we selected host genes of different circRNAs for Gene Ontology (GO) analysis and Kyoto Encyclopedia of Genes and Genomes (KEGG) analysis. The circRNAs were classified into three categories of the GO database: biological processes, cellular components and molecular functions. KEGG database was used to ascribe identified circRNAs to particular biological mechanisms and cellular pathways. (the established criteria: p adjusted < 0.05). GO and KEGG enrichment analysis was performed using (http://geneontology.org and http://www.kegg.jp/kegg).

### Microinjection

Three *circARMC4* siRNA species were designed to target different sites of porcine *circARMC4* sequence (GenePharma, Shanghai, China). All siRNA sequences used in the present study are shown in Supplementary Table S1. Microinjection was performed in T2 (TCM199 with 2% FBS) medium with 7.5 μg/ml CB on the heating stage of an inverted microscope (Olympus, Japan). Approximately 10 pl siRNA solution (20 μM) was microinjected into the cytoplasm of denuded GV oocytes. Two control groups (uninjected and RNase free water injection) were designed to exclude potential interferences of the microinjection technique. Oocytes from three groups were then matured *in vitro* for 44 h.

### Parthenogenetic activation

MII oocytes were stimulated with two pulses of direct current (1.56 kV/cm for 80 ms) by Cell Fusion Instrument (CF-150B, BLS, Hungary) in a chamber covered with activation medium (0.3 M mannitol supplemented with 0.1 mM CaCl_2_, 0.1 mM MgCl_2_ and 0.01% polyvinyl alcohol). Embryos were then incubated for 4 h in the chemically assisted activation medium (PZM-3 supplemented with 10 μg/ml cycloheximide and 10 μg/ml cytochalasin B). Next, embryos were cultured in fresh PZM-3 medium at 38.5 °C, 5% CO_2_, and 95% air with saturated humidity.

### Real-time quantitative PCR

Cumulus cells and oocytes were collected at GV and MII stage, respectively. Total RNA was extracted according to the manual of RNeasy Micro Kit (Qiagen) and was then incubated for 30 min at 37 °C with or without 5 U μg ^−1^ of RNase R (Epicentre Bio-technologies). Reverse transcription was performed using a QuantiTect Reverse Transcription Kit (Qiagen, 205311) according to the manufacturer’s instructions. Quantitative PCR was conducted using the SYBR Green PCR Master Mix (Roche, 04673514001) on a StepOne Plus Real-Time PCR System (Applied Biosystems). Reactions were carried out under the following conditions: 1 cycle of 95 °C for 2 min and 40 cycles of 95 °C for 5 s, 60 °C for 10 s. Analysis of gene expression employed relative quantification and 2 -ΔΔCT method. *EF1α1* was used as internal control. The PCR products were then run on 1.5 % agarose gels. The predicted strands were cut out directly for Sanger sequencing. Quantification of the fold change in gene expression was calculated using the comparative Ct (2^−ΔΔCt^) method. All the primers used are shown in Table S1. Three independent replicates were performed for each experiment.

### Immunofluorescence staining

Oocytes were fixed with 4% paraformaldehyde (PFA) for 20 min. After washing three times, the fixed oocytes were permeabilized with 1% Triton X-100 in DPBS for 30 min at room temperature (RT) and then blocked with 2% BSA in DPBS at RT for 1 h. Oocytes were incubated in blocking buffer containing α-Tubulin antibody (Sigma, F2168, 1:200) overnight at 4°C, After washing 3 times with DPBS for 60 min. Finally, oocytes were counterstained for 10 min in 4, 6-diamidino-2-phenylindole dihydrochloride (DAPI) solution and loaded onto glass slides followed by being covered with a glass coverslip. Samples were imaged using confocal microscopy (Olympus, Tokyo, Japan). At least 10 oocytes per group were used for each experiment. The specificity of the α-Tubulin antibody has been validated in Fig. S1.

### Statistical analysis

All experiments were carried out at least three times. The data were analyzed using either Student’s *t-*test or one-way ANOVA (SPSS 17.0) and were presented as mean ± standard error of mean (mean ± S.E.M). *P*<0.05 was considered to be statistically significant.

## Acknowledgments

We thank MS. Lulu Song for her help in technical assistance.

## Author contributions

Conceived and designed the experiments: ZBC YHZ. Performed the experiments: DG TTX LZ XT YQW DDZ WN XQ YYM KYJ TY. Analyzed the data: ZBC DG YSL YHZ. Contributed reagents and materials: ZBC YHZ. Wrote the paper: ZBC YHZ.

---

**Fig. S1.**
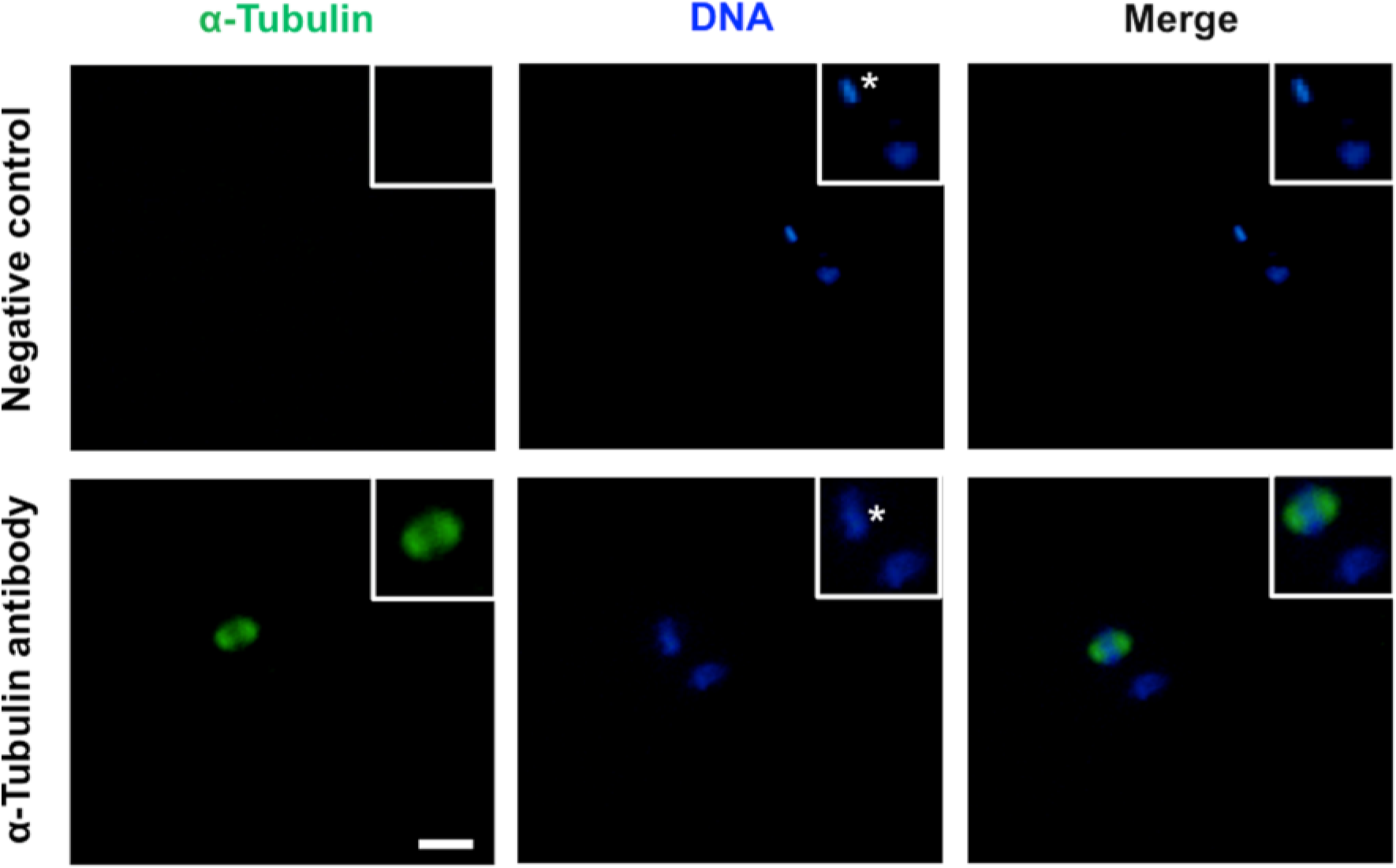
Validation of the specificity of α-tubulin antibody. MII oocytes were incubated with the diluent α-tubulin antibody in the immunofluorescence staining protocol while MII oocytes incubated with the mouse serum (Yeasen, 36118ES03) replacing α-tubulin antibody serve as a negative control. DNA was stained with DAPI (blue). Shown are representative images obtained using confocal laser-scanning microscopy. Right panel in each group shows the merged images between α-tubulin and DNA. White square insets indicate both spindles and chromosomes at high magnification. Asterisks mark chromosomes. Scale bar: 50 μm.

**Fig. S2.**
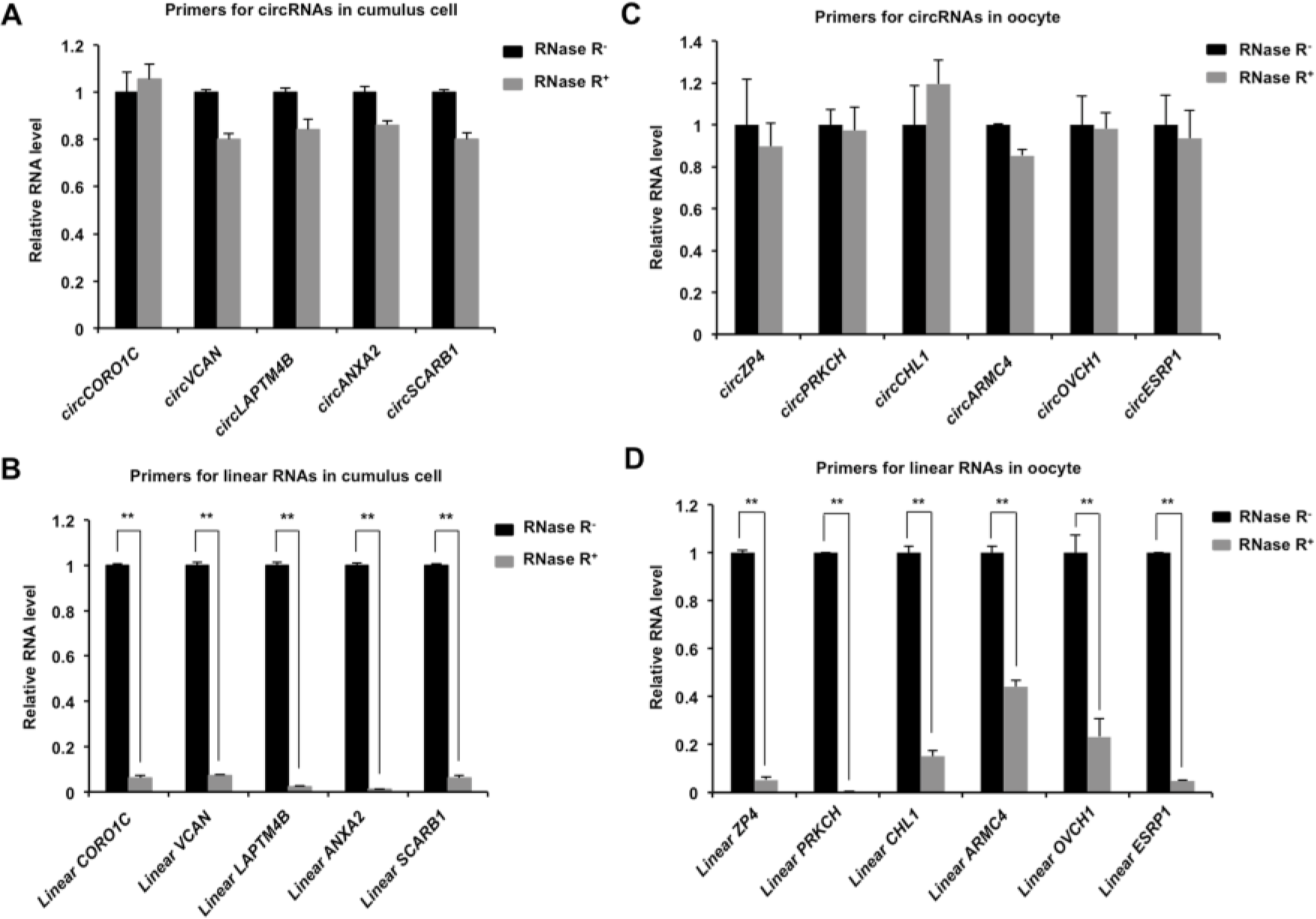
Expression of circRNAs and their corresponding linear RNAs in MCC and GV oocyte treated with or without RNase R. The several selected circRNAs and their corresponding linear RNAs were chosen from top differentially expressed circRNAs in both the cumulus cell and the oocyte. Total RNAs were extracted from MCC and GV oocytes, and then were treated with RNase R. The total RNAs without RNase R treatment serve as a control. Relative expression levels of the indicated circRNAs (A, C) and the corresponding linear RNAs (B, D) before and after RNase R treatment were determined by qPCR. The data were normalized against endogenous housekeeping gene *EF1α1*, and the value for both the MCC and the GV oocyte in the control group was set as one. The data are shown as mean ± S.E.M. Statistical analysis was performed using *t*-student test. Values with asterisks vary significantly, \*\**P* < 0.01.

